## Supplemental Tables for "SARS-CoV2, a threat to marine mammals? A study from Italian seawaters"

**Supplementary Material**

Fig. 1 SM: UWWTD Agglomerations - Big cities/dischargers by treatment type Map

European Environmental Agency thematic page (<https://www.eea.europa.eu/themes/water/european-waters/water-use-and-environmental-pressures/uwwtd>)

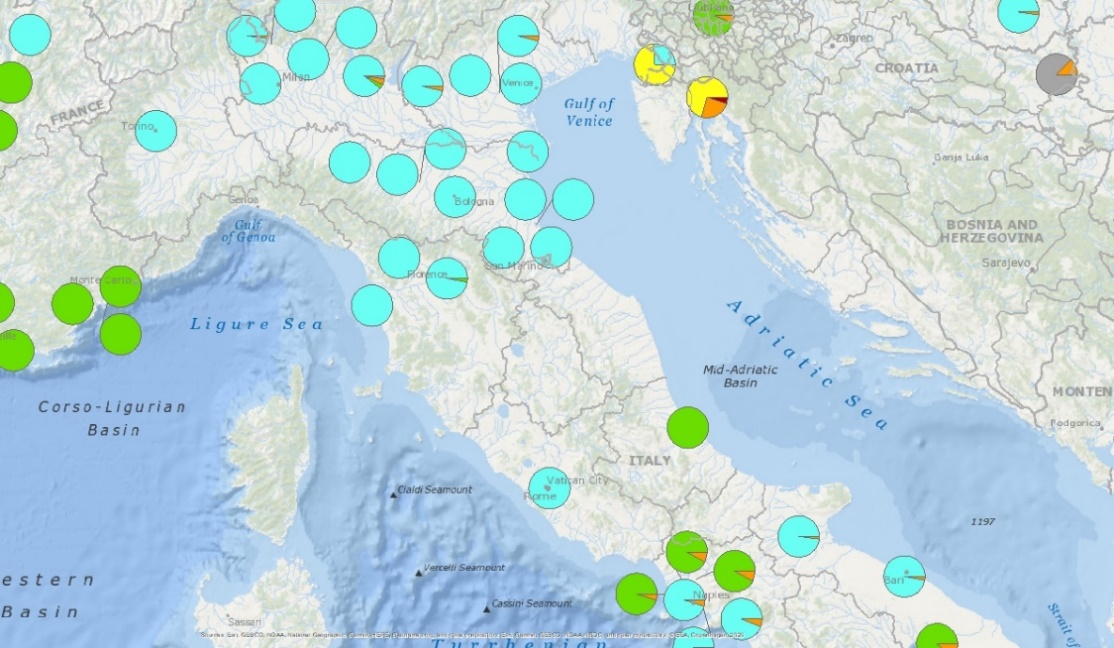

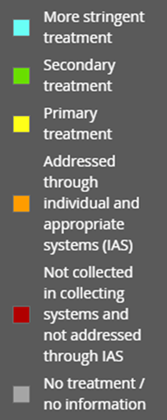

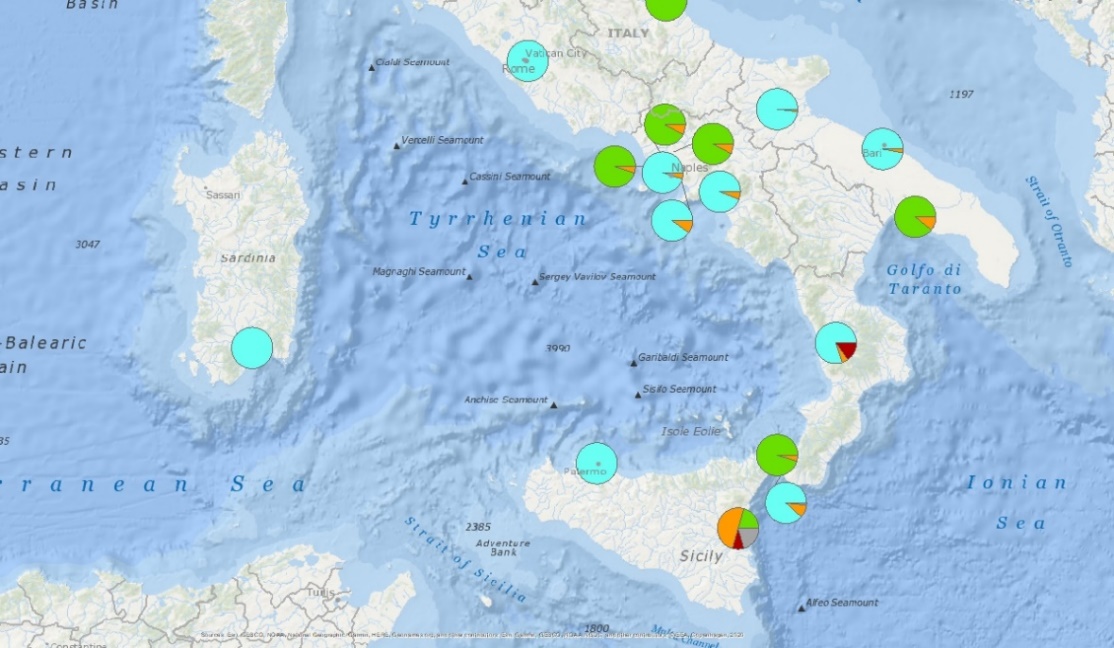

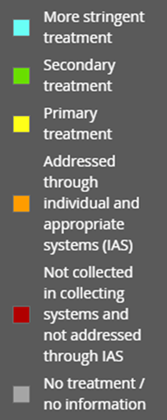

The size of each segment **within** the pie charts is proportional to the percentage of UWWTPs with specific treatment type**s**, where the total number of UWWTD is 100%. The segments are colored according to treatment type.

Fig. 2 SM: UWWTD plants by treatment type Map

European Environmental Agency thematic page (<https://www.eea.europa.eu/themes/water/european-waters/water-use-and-environmental-pressures/uwwtd>)

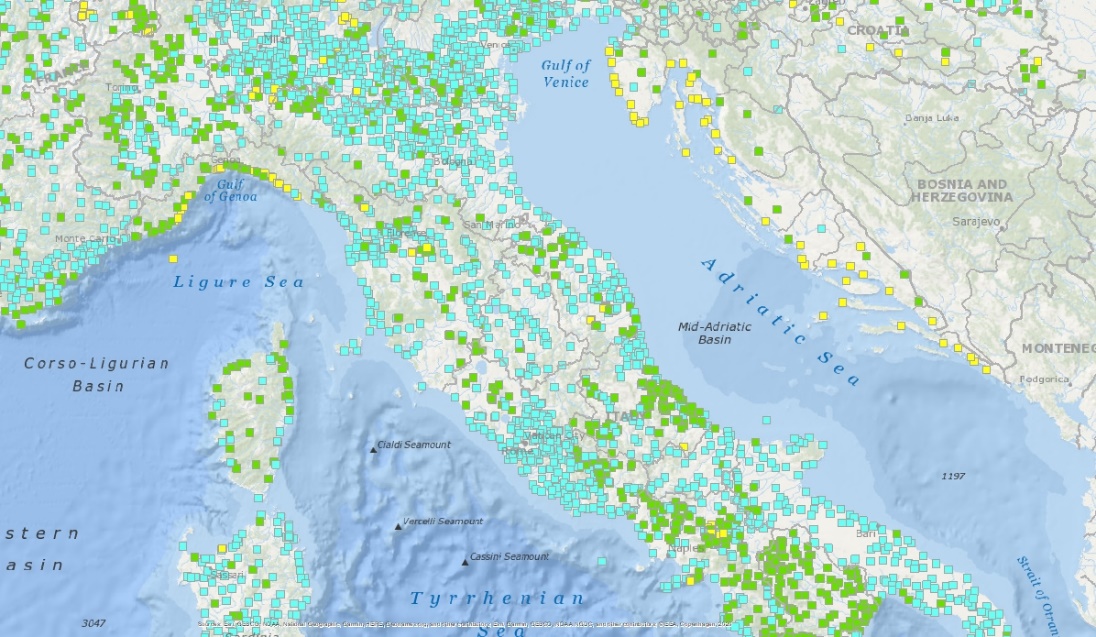

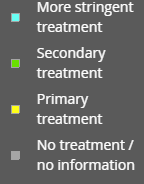

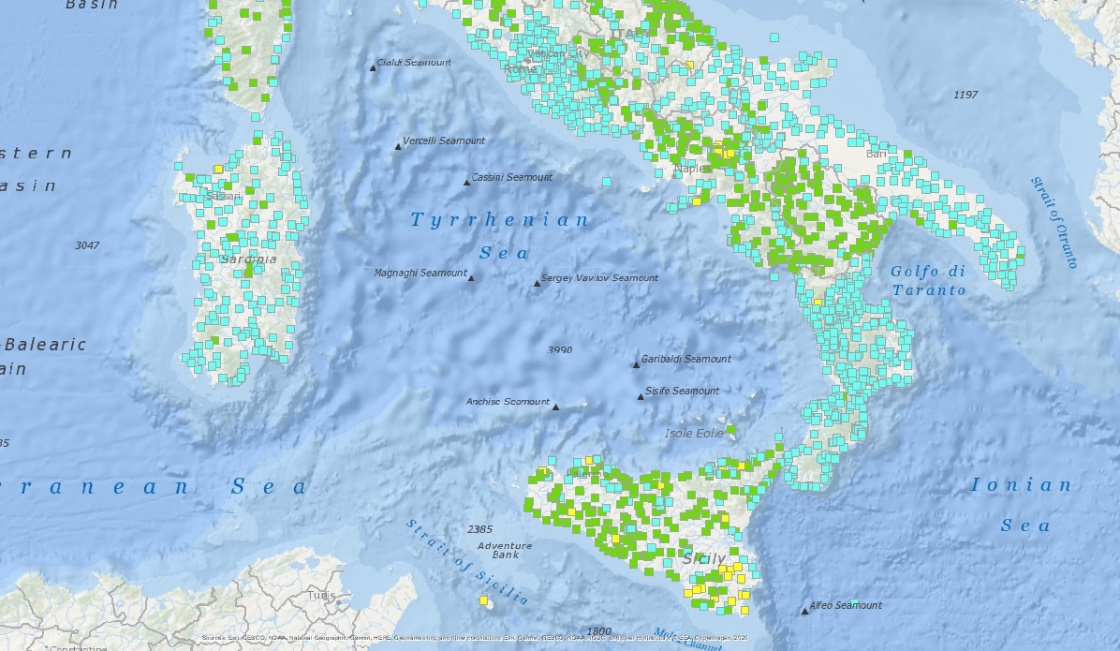

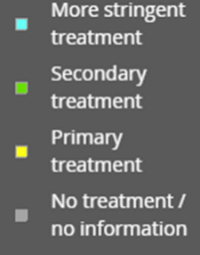

The dots representing the plants in the map are coloured according to treatment type.

Fig. 3 SM: UWWTD plants equipped with additional polishing treatment steps Map

European Environmental Agency thematic page environmental-pressures/uwwtd)

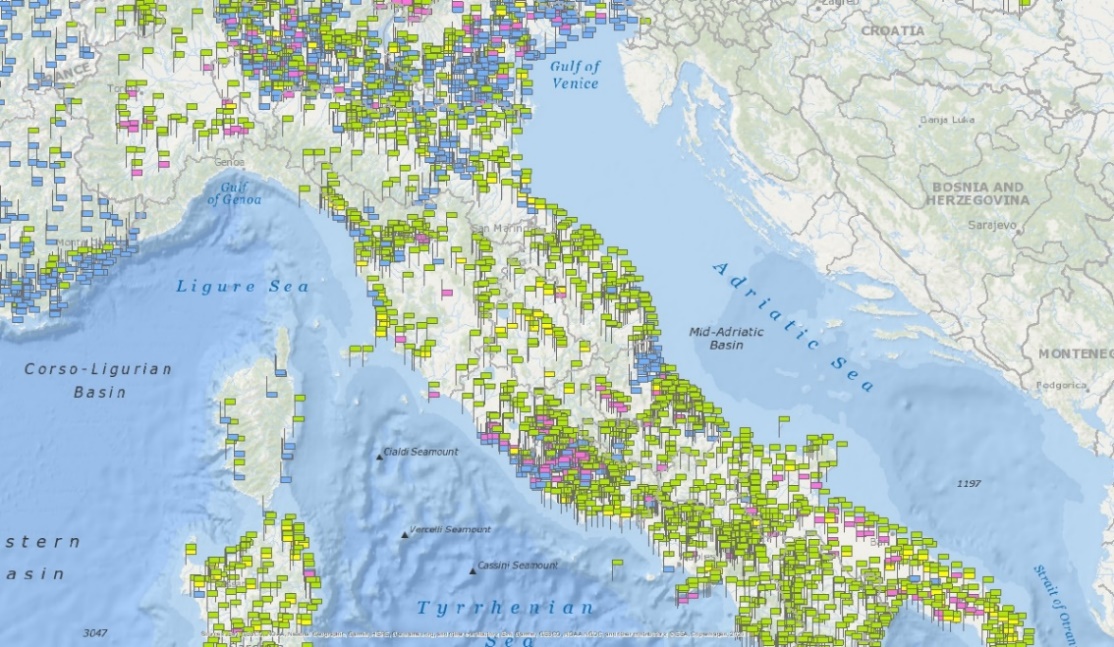

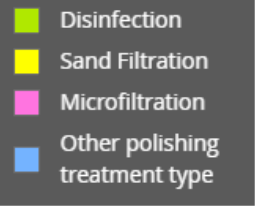

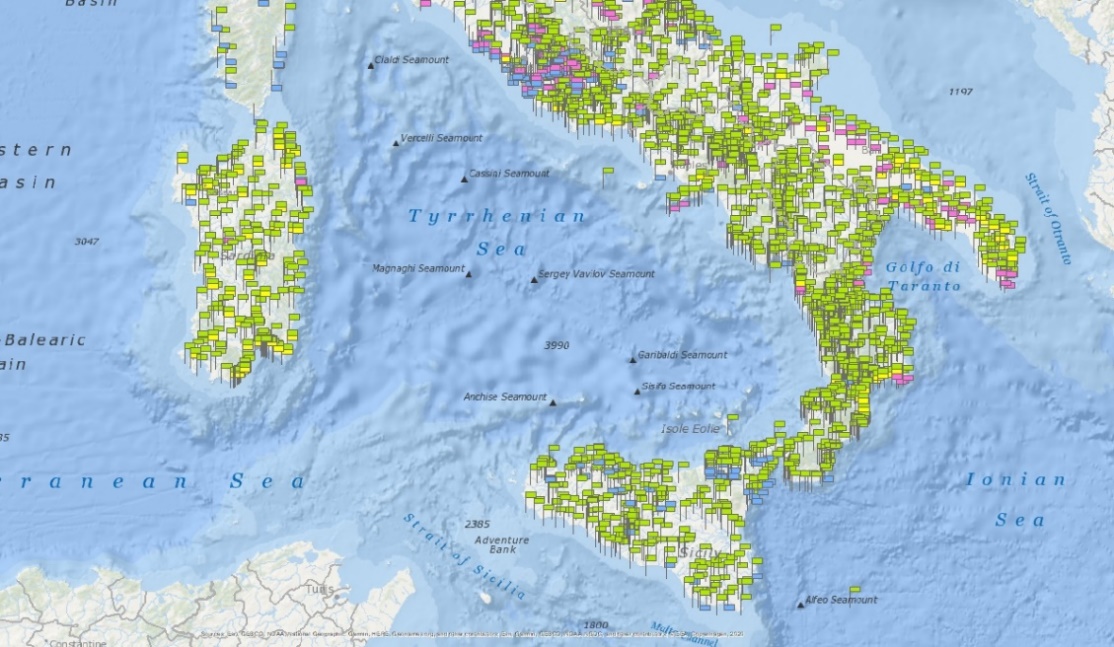

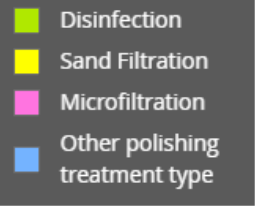

The dots representing the plants in the map are coloured according to treatment type.

Fig. 4 SM: Italian hydrogeological risk’s map.

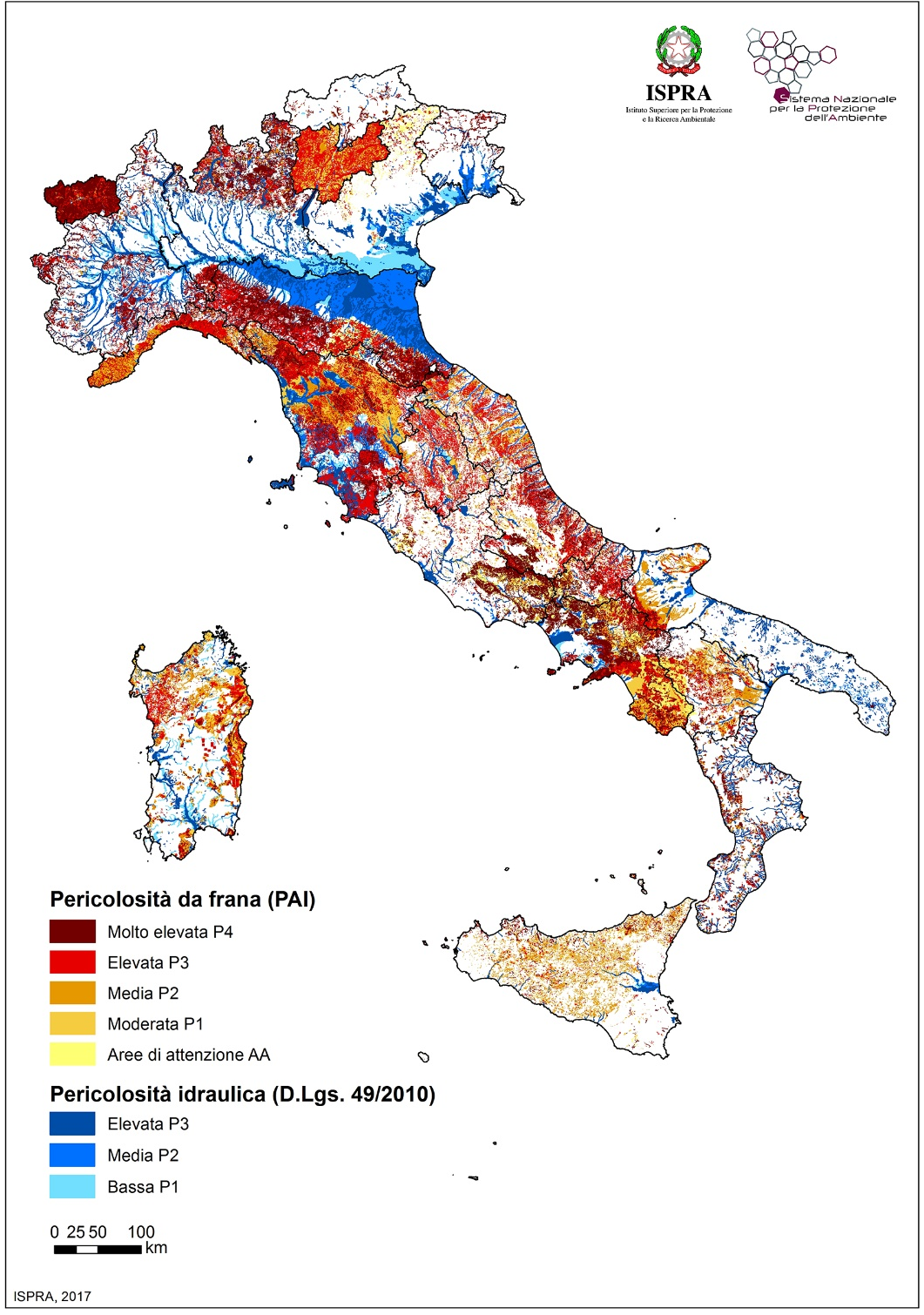
